# AB-Free Kava Reduces Anxiety-Like Behavior Without Preventing Nicotine-Induced Exploration Suppression in Mice

**DOI:** 10.64898/2026.08.11.744299

**Authors:** Guido Huisman, Lara S Caglayan, Marcelo Febo, Tengfei Bian, Yifan Wang, Chengguo Xing, Adriaan W. Bruijnzeel

## Abstract

Tobacco use is the leading preventable cause of death worldwide. Anxiety increases the risk for smoking, and smoking in turn increases the risk for anxiety disorders. There is therefore a need to identify interventions that reduce anxiety, in general and in the context of smoking, without producing sedation. Kava (Piper methysticum), a natural product with a long history of indigenous use, has been shown to have anxiolytic and calming effects and reduce nicotine withdrawal. The current study examined whether kava without the hepatotoxic flavokavains A and B (AB-free) could reduce anxiety-like behavior in mice repeatedly treated with nicotine. Male and female C57BL/6NCrl mice received either a control diet or an AB-free kava-supplemented diet and underwent two blocks of nicotine treatments. Mice underwent a first block of five every-other-day injections of nicotine (0.5 mg/kg) or saline, with open field testing after each injection, followed one week later by a nicotine challenge. A second block of injections was given using the same injection schedule, followed by a second challenge one week later, and two weeks afterward mice received a final challenge in a novel open field. During the first treatment block, AB-free kava significantly increased center time overall, an effect most pronounced in saline-treated animals, and increased locomotor activity, while nicotine decreased both measures. During the second challenge, nicotine reduced center time but not locomotor activity, and AB-free kava increased center time in saline-treated animals only. During the final challenge, nicotine reduced both measures, whereas AB-free kava increased center time regardless of nicotine treatment, and kava-treated animals also showed a near-significant increase in center entries. These results suggest that AB-free kava reduces anxiety-like behavior without inducing sedation but does not prevent nicotine-induced suppression of exploratory behavior.

## 1 Introduction

Tobacco use remains one of the most significant preventable causes of death globally, affecting an estimated 1.2 billion smokers and killing more than 7 million people annually (WHO, 2025). Smoking contributes to numerous life-threatening conditions, including chronic obstructive pulmonary disease, cardiovascular disease, stroke, and cancer (Ambrose and Barua, 2004; Forey et al., 2011; Ott et al., 1998; Sasco et al., 2004). Nicotine, the primary psychoactive alkaloid in tobacco and vaping products, can improve cognitive performance, induce mild euphoria, and provide acute relief from stress, and these short-term effects are believed to be a key reason for people to start or continue smoking (Wang et al., 2023; Stolerman and Jarvis, 1995; Pomerleau and Pomerleau, 1992). However, it is well established that chronic nicotine use can worsen anxiety symptoms in the long-term (Parrott, 1998; Moylan et al., 2013). Mechanistically, nicotine activates the hypothalamic-pituitary-adrenal (HPA) axis and thereby causes the release of stress peptides and hormones such as corticotropin-releasing hormone (CRH), adrenocorticotropic hormone (ACTH), and cortisol (Matta et al., 1998). Moreover, chronic exposure to nicotine can functionally alter neurotransmitter systems that regulate anxiety, such as gamma-aminobutyric acid (GABA), glutamate, and dopamine circuits (Benowitz, 2009; Picciotto and Kenny, 2021). Meta-analyses have found that smokers have significantly greater odds of anxiety compared to non-smokers, and that smoking cessation is associated with reductions in anxiety (Taylor et al., 2014b; Wu et al., 2023; Taylor et al., 2014a). Paradoxically, individuals with anxiety disorders are more likely to use nicotine products, potentially as a form of self-medication to manage acute anxiety symptoms, creating a cycle of dependence that ultimately worsens anxiety over time (Morissette et al., 2007; Cougle et al., 2010).

Animal models have been developed to study the effects of nicotine on the brain. Some studies have shown that repeated nicotine administration in mice leads to sensitized locomotor responses as indicated by a progressive enhancement in locomotor activity (Honeycutt et al., 2020; Correll et al., 2009; Ur Rehman et al., 2020). It has been suggested that this reflects the underlying neuroadaptations in dopamine-related reward circuitry and contributes to the development of addiction (Mao and McGehee, 2010; Vezina et al., 2007). Furthermore, nicotine administration in mice has been shown to induce anxiety-like behavior, as indicated by increased anxiety-like responses in the light/dark box and elevated plus maze test (Ouagazzal et al., 1999; Biala and Budzynska, 2006). These findings suggest that nicotine administration leads to adaptations that contribute to the development of nicotine addiction and increased anxiety.

Anxiety disorders are the most common psychiatric disorders in the United States, and people often self-medicate with nicotine products to temporarily improve mood and reduce feelings of stress (Bruijnzeel, 2012). However, although people often perceive smoking as relaxing, nicotine can paradoxically contribute to increased anxiety over time (Parrott, 1998; Moylan et al., 2013). Therefore, there is an urgent need for exploring alternative approaches for managing stress and anxiety in both smokers and nonsmokers. One possible approach that is rooted in traditional practices is the use of kava. Kava is historically consumed as a beverage among South Pacific Islanders for its stress-reducing and sleep-enhancing properties (Bian et al., 2020; WHO, 2016). Kava is traditionally prepared by grinding the roots of *Piper methysticum* and mixing them with water. Kava is rich in bioactive compounds known as kavalactones, which are believed to modulate neurotransmitter systems associated with relaxation and stress and anxiety reduction, an effect supported by clinical trial data (Pittler and Ernst, 2000; Volz and Kieser, 1997; Kuchta et al., 2021).

Interestingly, the use of kava is often associated with tobacco smoking. A study with kava users in the Republic of Vanuatu in the South Pacific suggests that smoking occurs most frequently in kava bars rather than at home and that about a quarter of regular kava drinkers reported regular smoking, compared to only four percent of subjects who did not use kava (Olszowy et al., 2022). Similarly, a study of Tongan men living in Australia found that about a quarter of participants smoked during kava drinking sessions (Maneze et al., 2008). These studies highlight the association between kava consumption and smoking.

Several clinical trials suggest that standardized kava extracts may reduce anxiety symptoms and may have fewer cognitive side effects and lower dependence liability than some conventional anxiolytic medications (Baric et al., 2018; Boerner et al., 2003; Cagnacci et al., 2003; Kuchta et al., 2021; Malsch and Kieser, 2001). In the United States, a wide variety of kava-based products are currently marketed as “calming” dietary supplements, but these formulations vary widely in form, composition, and concentration (Bian et al., 2020). Kavalactones are the active compounds in kava and are responsible for kava’s anxiolytic, mood-enhancing, and relaxing effects (Boonen et al., 1998; Boonen and Haberlein, 1998; Dinh et al., 2001; Yuan et al., 2002; Uebelhack et al., 1998; Magura et al., 1997) Kava contains six primary kavalactones, which serve as key standards for product formulation and quality (Bian et al., 2020). Although kava has been consumed safely for centuries, reports of severe hepatotoxicity and liver failure surfaced in the late 1990s and early 2000s, and there remain some concerns regarding kava’s hepatotoxic risk. The hepatotoxicity may be associated with non-traditional extraction methods or the use of low-quality kava cultivars. Traditional kava is prepared as an aqueous suspension, whereas certain organic extracts contain elevated levels of flavokavains A and B, which are highly cytotoxic (Zhou et al., 2010; Côté et al., 2004). These flavokavains are also more abundant in low-quality kava cultivars (Lebot et al., 2014). To mitigate these risks, a specialized kava formulation has been developed that includes the six major kavalactones while excluding flavokavains A and B (named AB-free kava), thereby reducing the likelihood of adverse health effects (Kanumuri et al., 2022).

In our previous work, we demonstrated the potential of AB-free kava to reduce smoking and, in a mouse model, to reduce lung inflammation after smoke exposure, to alleviate somatic nicotine withdrawal signs and to potentially improve lung function in smoke-exposed mice (Bian et al., 2024; Wang et al., 2020). However, it is currently unknown whether AB-free kava affects locomotor activity and anxiety-like behavior in mice repeatedly exposed to nicotine. Therefore, in the present study, we evaluated the effects of dietary administration of AB-free kava on locomotor activity and anxiety-like behaviors in male and female mice. Additionally, since kava is often used in conjunction with tobacco products, we investigated whether AB-free kava affects the acute effects of nicotine in open field tests. Both males and females were included because there are significant sex differences in anxiety-like behavior and in the effects of nicotine (Tan et al., 2019; Knight et al., 2021; Kim and Picciotto, 2023).

## 2 Materials and Methods

### 2.1 Animals

Adult male (20–26 g, 8 weeks of age; N = 36) and female (17–21 g, 8 weeks of age; N = 32) C57BL/6NCrl mice were purchased from Charles River (Raleigh, NC). The mice of the same sex were housed in groups of four in a climate-controlled vivarium (22°C with 50% humidity) with a 12 h light-dark cycle (lights on between 07:00–19:00). Food and water were available ad libitum. The experimental protocols were approved by the University of Florida Institutional Animal Care and Use Committee (IACUC). All experiments were performed in accordance with relevant IACUC guidelines and regulations and in compliance with ARRIVE guidelines 2.0 (Animal Research: Reporting of *In Vivo* Experiments).

### 2.2 Experimental design

Following a 2-week acclimatization period with no experimental interventions, the default ACS diets were removed from all cages on day 1 (See Figure 1 for a schematic overview). Each cage was allocated to receive either the control diet (AIN-93M powdered) or the AB-free kava diet, with the latter consisting of AIN-93M powdered diet supplemented with AB-free kava at a dose of 0.75 mg per gram of diet as in our previous work (Wang et al., 2025; Bian et al., 2024). Male and female mice were randomly assigned to one of four experimental groups: Control-Saline, Control-Nicotine, Kava-Saline, and Kava-Nicotine. All cages were maintained on their respective diets for the remainder of the study.

**Figure 1.**
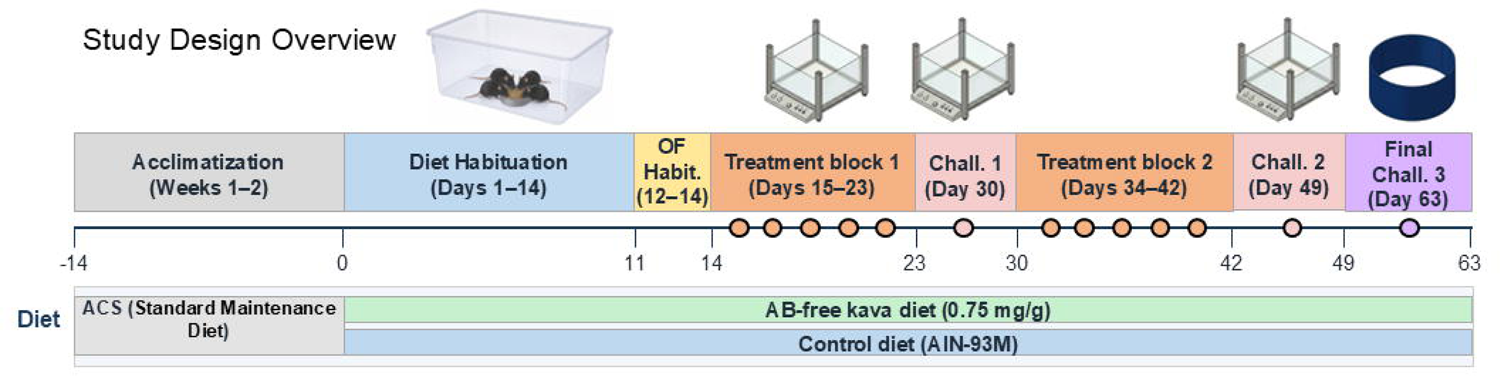
Schematic overview of the experimental timeline. Male and female mice were acclimatized for 2 weeks before being assigned to one of four experimental groups: Control-Saline, Control-Nicotine, Kava-Saline, or Kava-Nicotine. Mice were then acclimated to their assigned control or AB-free kava diet before open field habituation. Animals subsequently underwent two rounds of nicotine or saline treatment followed by challenge tests. Behavioral data were collected and analyzed during sensitization round 1, challenge 1, challenge 2, and the final challenge. Sensitization round 1 and the first two challenge tests were conducted in a square transparent open field, whereas the final challenge was conducted in a novel circular open field with dark blue walls and a white floor.

The diet habituation period lasted 14 days (days 1–14), during which mice were handled and weighed. All mice were habituated to the square transparent open field, hereafter referred to as the open field, for 3 days with 20-minute sessions each day, without injections. During all repeated treatment and challenge sessions the mice received 0.5 mg/kg nicotine (expressed as free base) or saline vehicle intraperitoneally (IP) in a volume of 10 ml/kg body weight. Beginning on day 15, mice underwent the first treatment round, with sessions every other day for a total of 5 sessions (days 15, 17, 19, 21, and 23). During each session, mice received IP injections of either nicotine or saline and were subsequently placed into the open field for 20 minutes. On day 30, one week after the final session, the first challenge was administered using the same protocol. The second treatment round began on day 34, repeating the every-other-day schedule (days 34, 36, 38, 40, and 42). During the second treatment round the mice were returned to the home cage after the injections. On day 49, one week after the final session of the second round, the second nicotine challenge was administered. On day 63, mice received a final challenge and were tested in a blue circular open field for 10 min, hereafter referred to as the novel open field. Following IP injection of nicotine or saline, mice were placed into the novel open field, and locomotor activity was recorded for 10 minutes. Testing occurred between 13:00 and 17:00. Arenas were cleaned with Nolvasan between mice.

A total of 68 mice (36 males and 32 females) were purchased. Prior to the start of the experiment, three male mice were lost due to fighting-related complications, resulting in a starting sample of 65 mice (33 males, 32 females; N = 8-9 per group). All 65 animals completed the sensitization procedures and the first two nicotine challenges. After completion of these procedures but prior to the novel open field test, two additional male mice were lost due to fighting-related complications. As a result, the sample size for the novel open field test (third challenge) was n = 63 (31 males, 32 females; N = 7-9 per group).

### 2.3 Drugs

(−)-Nicotine hydrogen tartrate salt (NIDA Drug Supply Program) was dissolved in sterile saline (0.9% sodium chloride), filtered through a 0.22-µm sterile filter, and adjusted to a pH of 7.4 ± 0.2 using 1 M NaOH. Nicotine was administered intraperitoneally at a dose of 0.5 mg/kg (expressed as free base) in a volume of 10 ml/kg body weight. Control animals received equivalent volumes of sterile saline.

### 2.4 Open field test

Locomotor activity and time in center were measured using an automated rodent activity monitoring system (AccuScan Instruments, Columbus, OH) (Chellian et al., 2021). The activity cages were constructed from clear Plexiglas (40 × 40 × 30 cm; L x W x H) with an outer steel frame equipped with infrared emitting and receiving panels. Each panel contained 16 evenly spaced infrared beams (2.5 cm apart) crossing the length and width of the cage at a height of 2 cm, enabling detection of horizontal movements. Patterns of sequential and repetitive infrared beam interruptions were recorded and transmitted to a programmable counter, which sorted the beam counts and sent the data to a PC running VersaMax™ software. The activity data were categorized into total distance traveled and time in the center of the cage (25 x 25 cm). Between each trial, the open field was cleaned with a Nolvasan (chlorhexidine diacetate) solution.

### 2.5 Circular open field test

The plastic circular open field measured 55 cm in diameter and 42 cm in height, with dark blue walls and a white floor to create a high-contrast environment for tracking (Bruijnzeel et al., 2011). The open field was designed to assess locomotor activity and anxiety-like behavior in rodents. Behavioral data were recorded and analyzed using Noldus Ethovision XT software (Wageningen, The Netherlands). For the zone analysis, the perimeter border zone was defined as 10 cm wide, and the inner zone comprised the remaining center. At the end of the study, total distance traveled, and time in the center were determined. Between each trial, the open field was cleaned with a Nolvasan (chlorhexidine diacetate) solution.

### 2.6 Statistics

Data were analyzed using IBM SPSS Statistics (version 30, Armonk, NY). All data are presented as mean ± standard error of the mean (SEM). Locomotor activity and time spent in the center during the first five-day treatment period were analyzed using a mixed repeated-measures analysis of variance (ANOVA) with time as the within-subjects factor and diet, treatment, and sex as between-subjects factors. Data from the nicotine challenge tests were analyzed using a three-way ANOVA with diet, treatment, and sex as between-subjects factors. Body weight was analysed using a mixed repeated-measures ANOVA with Time as the within-subjects factor (24 timepoints) and sex, diet (control vs. kava), and treatment (saline vs. nicotine) as between-subjects factors. Significant interactions were followed up with post hoc pairwise comparisons using Bonferroni correction, with the effect of one factor tested within each level of the other factor(s). To account for sex as a biological variable, all post hoc comparisons were conducted separately for males and females. For all analyses, statistical significance was defined as p < 0.05, and exact p-values are reported except where p < 0.001. Figures were generated using GraphPad Prism (version 11, San Diego, CA).

## 3 Results

### 3.1 Open field test

#### 3.1.1 Distance traveled during the first five days of nicotine treatment

The analysis of total distance traveled across the five-day treatment period revealed significant effects of time, diet, and nicotine treatment (Figure 2A, B). The repeated measures ANOVA showed that locomotor activity changed over time (F(4, 228) = 6.038, p < 0.001). There was also a main effect of diet, with kava-treated animals traveling greater distances than animals that received the control diet (F(1, 57) = 6.719, p = 0.012). Additionally, nicotine treatment led to a significant decrease in the total distance traveled (F(1, 57) = 10.077, p = 0.002). No significant main effect of sex was observed (F(1, 57) = 0.464, p = 0.498), and the diet × treatment interaction was not significant (F(1, 57) = 1.071, p = 0.305), indicating that the effect of the kava diet on locomotor activity did not depend on nicotine treatment. No other significant interactions were detected.

**Figure 2.**
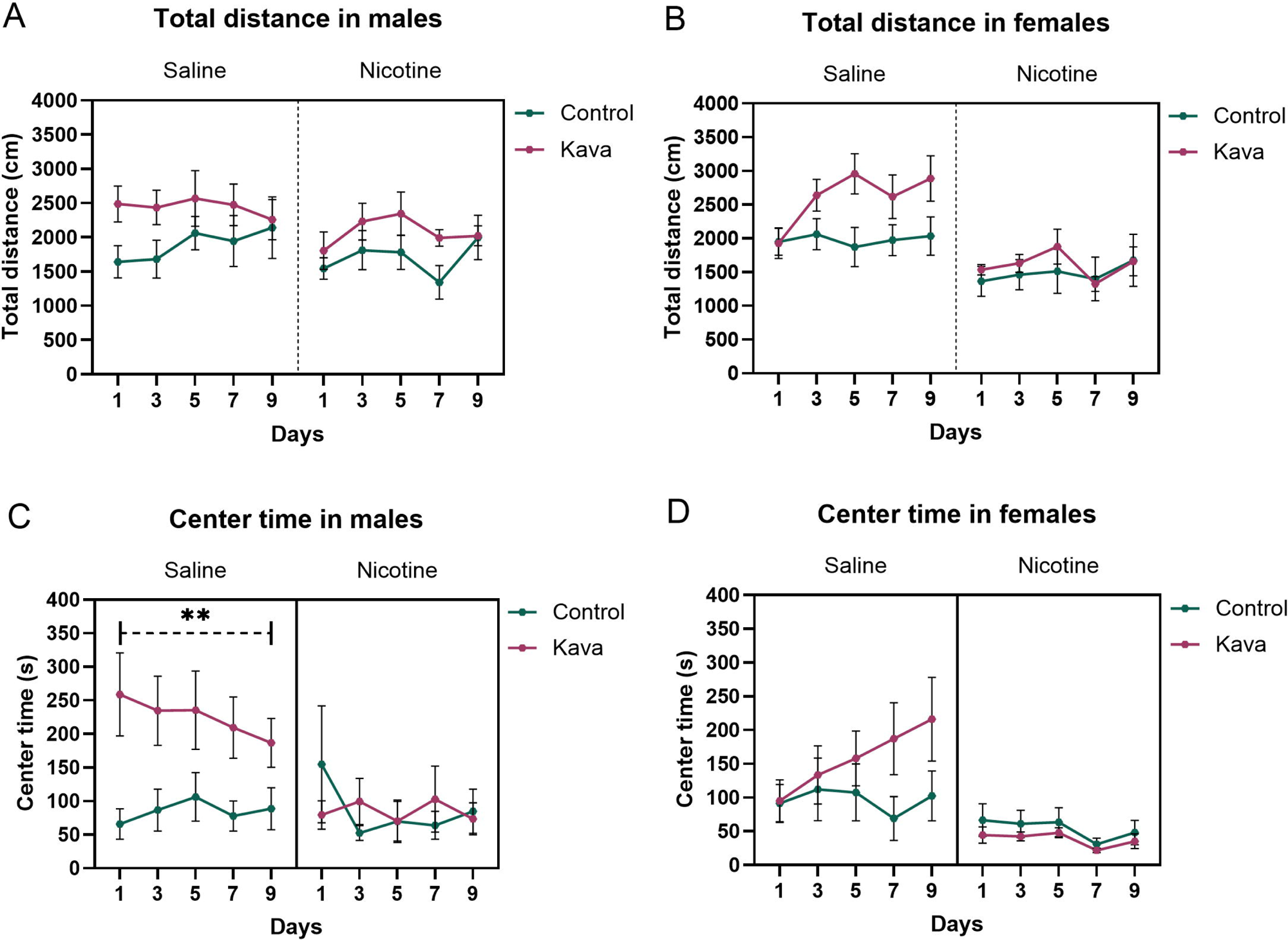
Locomotor activity and anxiety-like behavior in the open field over five days. Total distance traveled by male (A) and female (B) mice, and time spent in the center zone by male (C) and female (D) mice over five days. In panel (C), ** Kava-Saline males spent significantly more time in the center than Control-Saline males (p = 0.006). ** p < 0.01, Bonferroni corrected post hoc comparisons. N = 8-9 per group. Data are expressed as mean ± SEM.

#### 3.1.2 Time in center during the first five days of nicotine treatment

Analysis of time spent in the center of the open field over the five-day testing period revealed significant effects of treatment and a diet × treatment interaction (Figure 2C, D). Nicotine treatment significantly reduced the time in the center across groups (F(1, 57) = 11.680, p = 0.001). A significant main effect of diet was observed, with kava-treated animals spending more time in the center than control diet animals (F(1, 57) = 4.419, p = 0.040). There was also a significant diet × treatment interaction (F(1, 57) = 6.043, p = 0.017), indicating that the effect of the kava diet on center time depended on nicotine treatment. Bonferroni post hoc comparisons revealed that among males, Kava-Saline animals spent significantly more time in the center than Control-Saline animals (p = 0.006), whereas kava did not increase center time in nicotine-treated males (p = 0.998). No significant diet effects were observed in females under either condition (saline p = 0.155; nicotine p = 0.714).

### 3.2 First nicotine challenge

#### 3.2.1 Distance moved during first nicotine challenge

The effects of nicotine treatment, diet, and sex on locomotor activity were assessed (Figure 3A). Nicotine treatment significantly decreased the distance traveled in the open field (F(1, 57) = 8.666, p = 0.005). A significant three-way interaction of diet × treatment × sex was observed (F(1, 57) = 4.790, p = 0.033), indicating that the effects of the kava diet and nicotine treatment on locomotor activity differed between males and females. Post hoc Bonferroni comparisons in females revealed that kava-treated animals traveled significantly greater distances than Control-treated animals under saline conditions (p = 0.005) but not under nicotine conditions (p = 0.776).

**Figure 3.**
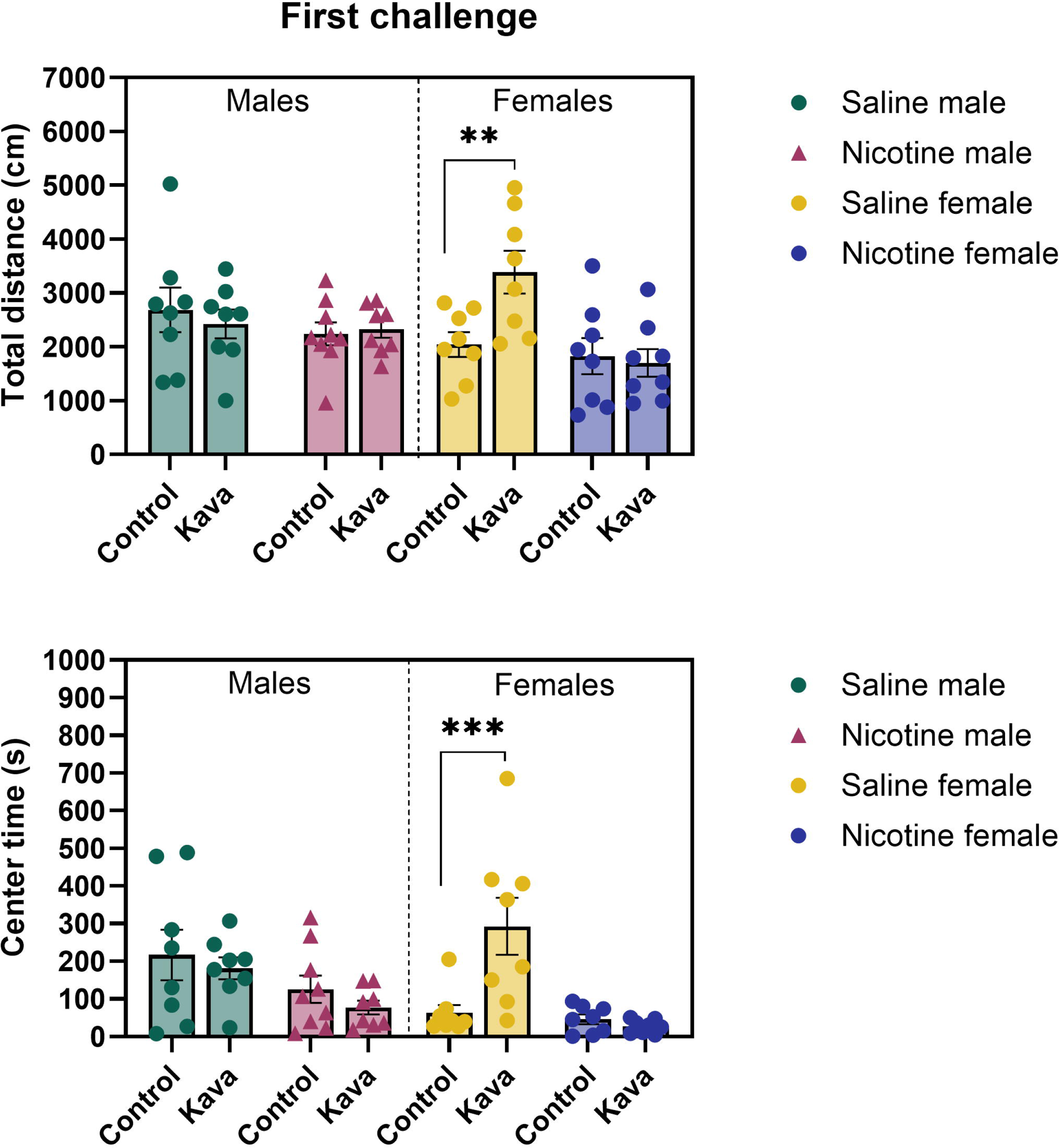
Locomotor activity and anxiety-like behavior in the open field during the first nicotine challenge. Total distance traveled by all eight experimental groups is shown in panel (A), while time spent in the center zone is depicted in panel (B). In panel (A), ** Kava-Saline females traveled significantly greater distances than Control-Saline females (p = 0.005). In panel (B), *** Kava-Saline females spent significantly more time in the center than Control-Saline females (p < 0.001). ** p < 0.01, *** p < 0.001. Bonferroni corrected post hoc comparisons. N = 8-9 per group. Data are expressed as mean ± SEM.

#### 3.2.2 Time in center during first nicotine challenge

Nicotine treatment significantly reduced center time across groups (F(1, 57) = 17.019, p < 0.001) (Figure 3B). There was a significant diet × sex interaction (F(1, 57) = 6.966, p = 0.011) and a significant three-way diet × treatment × sex interaction (F(1, 57) = 4.048, p = 0.049), indicating that the effects of the kava diet and nicotine treatment on center time differed between males and females. Post hoc Bonferroni comparisons revealed that among females, kava-treated animals spent significantly more time in the center than Control-treated animals under saline conditions (p < 0.001) but not under nicotine conditions (p = 0.740). No significant effects of diet were observed in males under either condition (saline p = 0.547; nicotine p = 0.344).

### 3.3 Second nicotine challenge

#### 3.3.1 Distance moved during the second nicotine challenge

The effects of nicotine treatment, diet, and sex on locomotor activity were assessed (Figure 4A). No significant main effects of nicotine treatment, diet, or sex were observed on distance traveled. However, a significant diet × treatment interaction was detected (F(1, 57) = 4.605, p = 0.036), indicating that the effect of the kava diet on locomotor activity depended on nicotine treatment. Post hoc Bonferroni comparisons revealed no significant diet effects in males under either saline or nicotine conditions (saline p = 0.277; nicotine p = 0.982). No significant diet effects were observed in females under either condition (saline p = 0.307; nicotine p = 0.065), although a trend was observed under nicotine conditions (p = 0.065).

**Figure 4.**
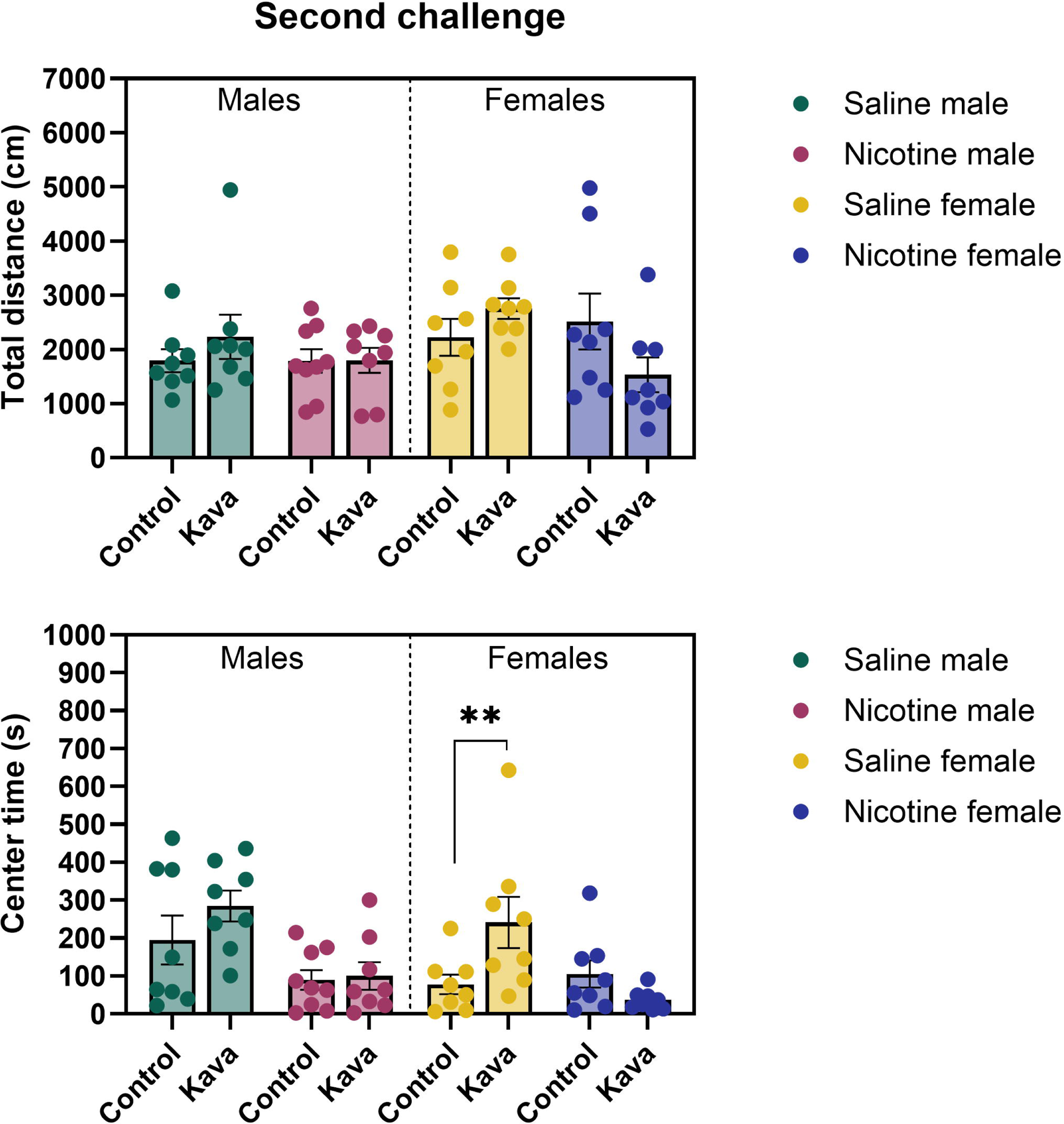
Locomotor activity and anxiety-like behavior in the open field during the second nicotine challenge. Total distance traveled by all eight experimental groups is shown in panel (A), while time spent in the center zone is depicted in panel (B). In panel (B), ** Kava-Saline females spent significantly more time in the center than Control-Saline females (p = 0.008). ** p < 0.01. Bonferroni corrected post hoc comparisons. N = 8-9 per group. Data are expressed as mean ± SEM.

#### 3.3.2 Time in center during the second nicotine challenge

Nicotine treatment significantly reduced center time across groups (F(1, 57) = 15.459, p < 0.001) (Figure 4B). A significant diet × treatment interaction was observed (F(1, 57) = 6.836, p = 0.011), indicating that the effect of the kava diet on center time depended on nicotine treatment. No significant main effects of diet or sex were observed, and no other significant interactions were detected. Post hoc Bonferroni comparisons revealed that among females, kava-treated animals spent significantly more time in the center than Control-treated animals under saline conditions (p = 0.008) but not under nicotine conditions (p = 0.243). No significant diet effects were observed in males under either condition (saline p = 0.160; nicotine p = 0.861).

### 3.4 Final nicotine challenge

#### 3.4.1 Distance moved during the third nicotine challenge

Analysis of distance moved in the novel open field revealed a significant main effect of nicotine treatment, with nicotine-treated animals traveling less than saline controls (F(1, 55) = 37.193, p < 0.001) (Figure 5A). No significant main effects of diet or sex were observed, and no significant interactions were detected.

**Figure 5.**
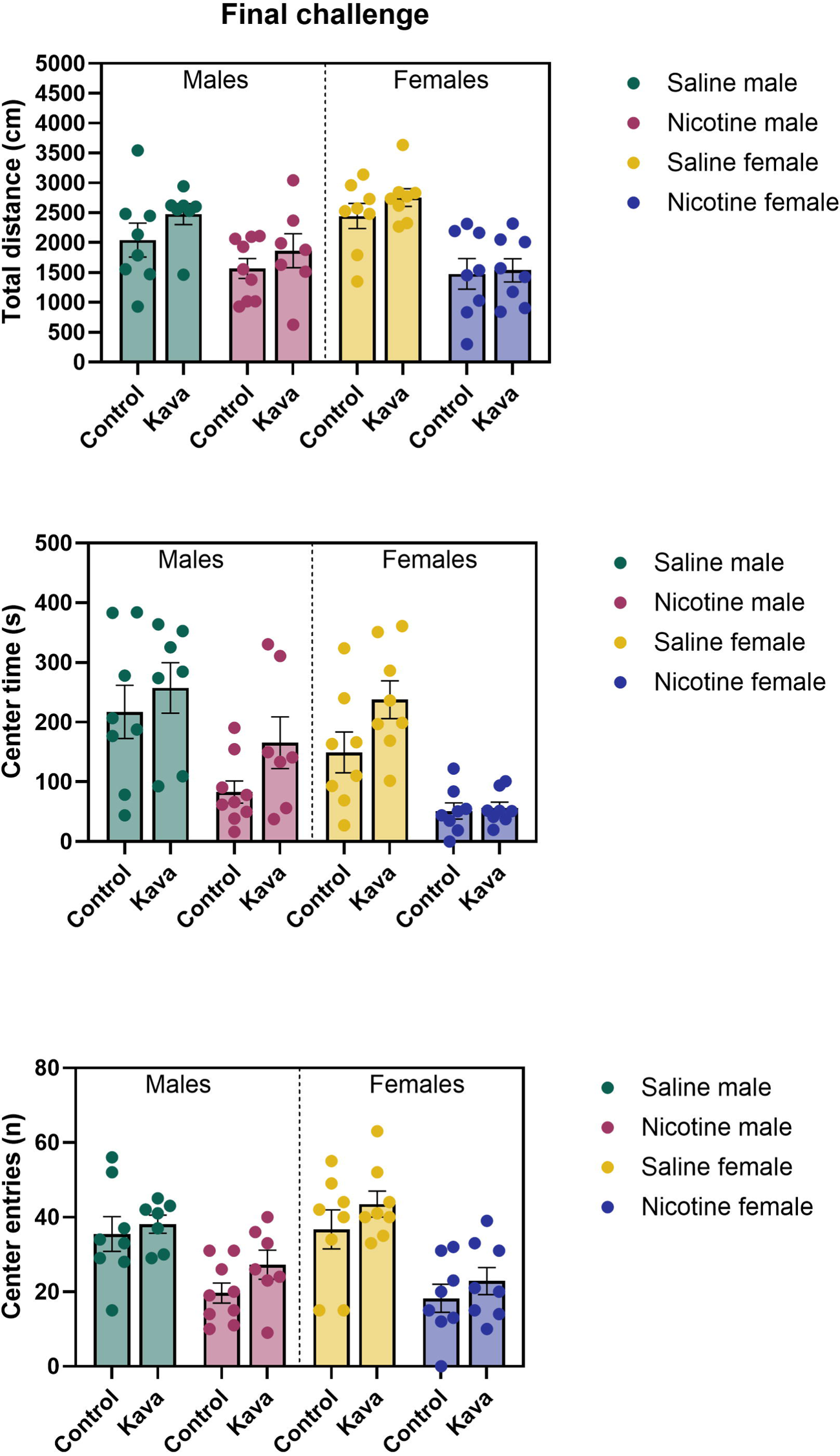
Locomotor activity and anxiety-like behavior in the novel open field during the final nicotine challenge. Total distance traveled by all eight experimental groups is shown in panel (A), time spent in the center zone in panel (B), and center entries in panel (C). N = 7-9 per group. Data are expressed as mean ± SEM.

#### 3.4.2 Time in center during third nicotine challenge

Analysis of time spent in the center of the novel open field revealed significant main effects of diet, treatment, and sex (Figure 5B). Kava-treated animals spent more time in the center compared to control diet animals (F(1, 55) = 5.892, p = 0.019). Nicotine treatment significantly reduced center time across groups (F(1, 55) = 32.232, p < 0.001). Additionally, males spent more time in the center than females (F(1, 55) = 6.635, p = 0.013). No significant interactions were observed.

#### 3.4.3 Center frequency during third nicotine challenge

Analysis of the number of entries into the center of the novel open field revealed significant main effects of nicotine treatment (Figure 5C). Nicotine-treated animals entered the center less frequently than saline controls (F(1, 55) = 36.716, p < 0.001). The main effect of diet approached but did not reach significance, with kava-treated animals tending to enter the center more frequently than control diet animals (F(1, 55) = 3.968, p = 0.051). No significant main effect of sex was observed (F(1, 55) = 0.005, p = 0.943), and no significant interactions were detected.

### 3.5 Body weights

There was a significant main effect of time (F(23, 1311) = 115.52, p < 0.001), indicating that body weight increased across the study period. There was a significant effect of sex on body weight (F(1, 57) = 405.80, p < 0.001), indicating that the males had a higher body weight compared to the females. The time × sex interaction was also significant (F(23, 1311) = 9.93, p < 0.001), indicating that males gained weight at a faster rate than females over the course of the study. Neither the kava diet nor nicotine treatment had a significant effect on body weight over the course of the experiment (see Supplementary Figure S2).

An additional analysis was conducted in which the body weights were expressed as a percentage of the pre-diet baseline (last measurement before the onset of the new diets; Figure S3). The analysis of the normalized data revealed no significant effect of diet (F(1, 57) = 2.637, p = 0.110) or nicotine treatment (F(1, 57) = 0.090, p = 0.766), confirming that neither the kava diet nor nicotine treatment significantly altered body weight gain relative to baseline. A significant effect of sex was observed (F(1, 57) = 4.647, p = 0.035), with males showing a greater relative weight gain than females. There was also a significant main effect of time (F(17, 969) = 98.177, p < 0.001), reflecting the increase in body weight over the course of the study, and a significant time × sex interaction (F(17, 969) = 6.431, p < 0.001), indicating that the trajectory of relative weight gain differed between males and females. A significant time × diet × sex interaction was also observed (F(17, 969) = 2.397, p = 0.001). However, Bonferroni-corrected post hoc comparisons revealed no significant differences between the kava and control diets at any individual timepoint in either sex.

## 4 Discussion

In the present study we investigated the effects of AB-free kava and repeated nicotine treatment on locomotor activity and anxiety-like behavior in male and female mice. Our study revealed significant effects of the AB-free kava diet, nicotine treatment, and sex on locomotor activity and center time in two different open field tests. During the initial five-day nicotine treatment period, AB-free kava increased locomotor activity, while nicotine reduced activity in all groups. AB-free kava also increased the time spent in the center of the open field, which is indicative of an anxiolytic-like effect. The effect of kava on center time was most pronounced in the saline-treated males. In contrast, nicotine reduced time in the center of the open field. These findings indicate that kava had anxiolytic effects in saline-treated animals but was less effective in animals exposed to nicotine. In the challenge tests, AB-free kava increased center time in saline-treated animals, whereas its effects on locomotor activity were less consistent. AB-free kava had limited efficacy in counteracting the anxiogenic effects of nicotine. The results of the novel open field test confirmed that AB-free kava increased center time, supporting an anxiolytic-like effect. Furthermore, nicotine decreased locomotor activity and center time across groups, and males spent more time in the center than females. These findings indicate that AB-free kava has anxiolytic-like effects, with mild and variable effects on locomotor activity, while nicotine has locomotor-suppressing and anxiogenic-like effects.

Our study examined whether repeated treatment with 0.5 mg/kg nicotine affected locomotor activity and anxiety-like behavior in male and female C57BL/6 mice. The repeated nicotine treatment schedule was based on established nicotine sensitization procedures (Honeycutt et al., 2020; Correll et al., 2009). Contrary to some previous studies reporting nicotine-induced sensitization of locomotor responses, we did not observe an increase in locomotor activity over time. Several studies have reported nicotine-induced locomotor sensitization under similar but not identical conditions. In one study, adult male and female C57BL/6 mice received five IP injections of nicotine (0.5 mg/kg) to examine nicotine sensitization (Honeycutt et al., 2020). On day 5, both male and female nicotine-treated mice demonstrated significantly higher activity levels compared to their saline-treated counterparts. In another study, adolescent male C57BL/6 mice were treated with subcutaneous (SC) nicotine (0.5 mg/kg free base) daily for 7 days (Correll et al., 2009). Following a one-week drug-free period, the mice received a challenge injection at the same dose. During the initial 7-day treatment period, nicotine reduced locomotor activity but after the challenge injection locomotor activity was increased compared to the saline group. In a related study, male BALB/c mice received 0.5 mg/kg nicotine IP for seven consecutive days and on a challenge day after a three-day drug-free period (Ur Rehman et al., 2020). Nicotine increased locomotor activity compared to saline controls on Day 7 of treatment and on the challenge day. However, it is important to note that not all studies report an increased locomotor response following repeated nicotine treatment. In another study, male C57BL/6NCrl mice (same substrain as in our study) received SC nicotine injections (0.5 or 1.0 mg/kg, base) for five days (Akinola et al., 2019). After a two-day drug free period, the mice were challenged with the same doses of nicotine. The lowest nicotine dose had no effect on locomotor activity while the higher nicotine dose decreased locomotor activity. Given that our study also used the same C57BL/6NCrl substrain, it is likely that we did not observe a sensitized locomotor response to nicotine because this substrain may not develop locomotor sensitization under standard sensitization protocols. In contrast, other strains, such as Swiss mouse strains, appear to develop locomotor sensitization across a wide range of nicotine doses (Ulusu et al., 2005; Biala and Staniak, 2010).

In the present study we also found that nicotine increased anxiety-like behavior in the open field tests as indicated by a decrease in center time. The effects of acute nicotine administration on anxiety-like behavior have been investigated before and studies have reported both anxiogenic and anxiolytic effects of nicotine, depending on factors such as dosage, timing, and the specific behavioral assays employed. However, few studies have investigated the effects of nicotine on anxiety-like behavior in the open field test. Notably, one study reported that administration of nicotine at 0.5 mg/kg (free base) in male and female Swiss mice did not affect anxiety-like behavior in the open field test (Dutra-Tavares et al., 2023). The elevated plus maze is one of the most widely used tests to investigate the effects of drug on anxiety-like behavior in rodents. In the mouse elevated plus maze test, acute nicotine administration has yielded varying outcomes. Nicotine (0.1–0.5 mg/kg) has been reported to produce anxiolytic effects (Brioni et al., 1993), anxiogenic effects (Ouagazzal et al., 1999; Biala and Budzynska, 2006), or no effects (Benwell et al., 1994). Studies with the social interaction test indicate that the time of nicotine administration affects anxiety-like behavior as nicotine has anxiogenic effect 5 min after administration but an anxiolytic effect 30 min after administration (Irvine et al., 1999). Our study shows that 0.5 mg/kg of nicotine has anxiogenic effects in different open field tests in male and female C57BL/6NCrl mice. However, as indicated above factors such as the mouse strain and testing conditions can affect the outcome of these studies.

In the present study, we found that AB-free kava reduced anxiety-like behavior as indicated by an increase in center time in the open field tests in male and female mice. However, kava also increased locomotor activity during the first treatment block, which raises the possibility that the concurrent increase in center time reflected greater overall activity rather than a specific reduction in anxiety-like behavior. Two observations argue against this interpretation. First, in the novel open field during the final challenge, kava significantly increased center time (p = 0.019) without affecting distance traveled, indicating that the two measures can dissociate. Second, kava-treated animals also showed a near-significant increase in the number of center entries during the final challenge (p = 0.051), which is a measure that is less directly tied to total distance than center duration (Prut and Belzung, 2003). Together, these findings suggest that the increased center time in kava-treated animals reflects reduced anxiety-like behavior rather than nonspecific hyperactivity. The anxiolytic properties of kava have also been demonstrated in several preclinical models. Consistent with our findings, Garrett et al. reported dose-dependent anxiolytic-like behavioral changes with a kava extract in the mirrored chamber avoidance assay and elevated plus maze in BALB/c mice (Garrett et al., 2003). Similarly, another study showed that a kava preparation (Kava-Kava extract LI 150) increased open-arm exploration in the elevated plus maze test in male Wistar rats, though its effects were less pronounced than those of diazepam (Rex et al., 2002). On a related note, anxiolytic-like effects of a kava extract and its active component, dihydrokavain, have been demonstrated in a chick social separation-stress paradigm (Smith et al., 2001). The present study advances the field by providing evidence that a standardized AB-free kava formulation, designed with an enhanced safety profile, effectively reduces anxiety-like behavior in adult mice. These findings underscore the potential of AB-free kava as a safe and effective alternative for managing anxiety disorders.

Overall, our findings provide insight into the anxiolytic effects of AB-free kava as well as the complex interplay between AB-free kava, nicotine, and sex in modulating locomotor activity and anxiety-like behavior. In our studies, AB-free kava generally increased locomotor activity and center time suggesting anxiolytic-like effects. However, these effects were attenuated in the presence of nicotine. Together, these findings support the potential of AB-free kava as an anxiolytic-like intervention.

### Funding Sources

Adriaan Bruijnzeel and Chengguo Xing were supported by Grant 23B02 from the Florida Department of Health.

## Acknowledgments

During the preparation of this manuscript, the authors used Claude Opus 4.8 (Anthropic) to improve the spelling, grammar, and flow of the text and to assist with preparing the supplementary figures. The authors reviewed and edited all content and take full responsibility for the accuracy and integrity of the manuscript.

## Notes

### Competing Interest Statement

The authors have declared no competing interest.

