## Supplement for "AB-Free Kava Reduces Anxiety-Like Behavior Without Preventing Nicotine-Induced Exploration Suppression in Mice"

Supplemental Figures

A

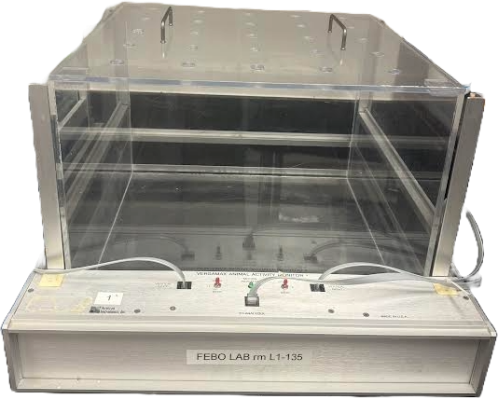

B

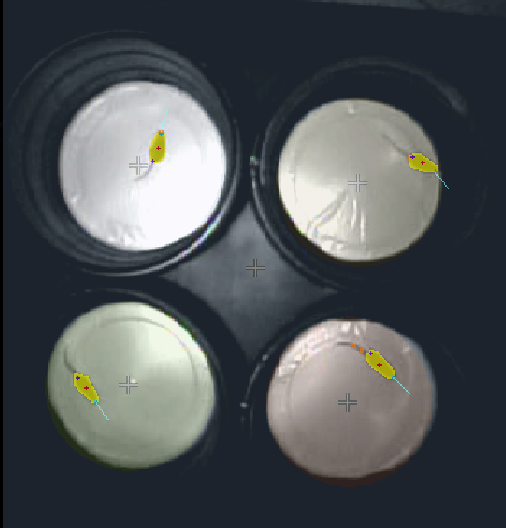

**Figure S1. Apparatus used for open field testing.** (A) AccuScan open field apparatus. Clear Plexiglas activity cages (40 × 40 × 30 cm; L × W × H) equipped with infrared beam panels for automated detection of locomotor activity and time in center (VersaMax™ software). (B) Circular open field arena (55 cm diameter, 42 cm height) with a white floor and dark blue walls. Four arenas shown simultaneously tracked using Noldus EthoVision XT software.

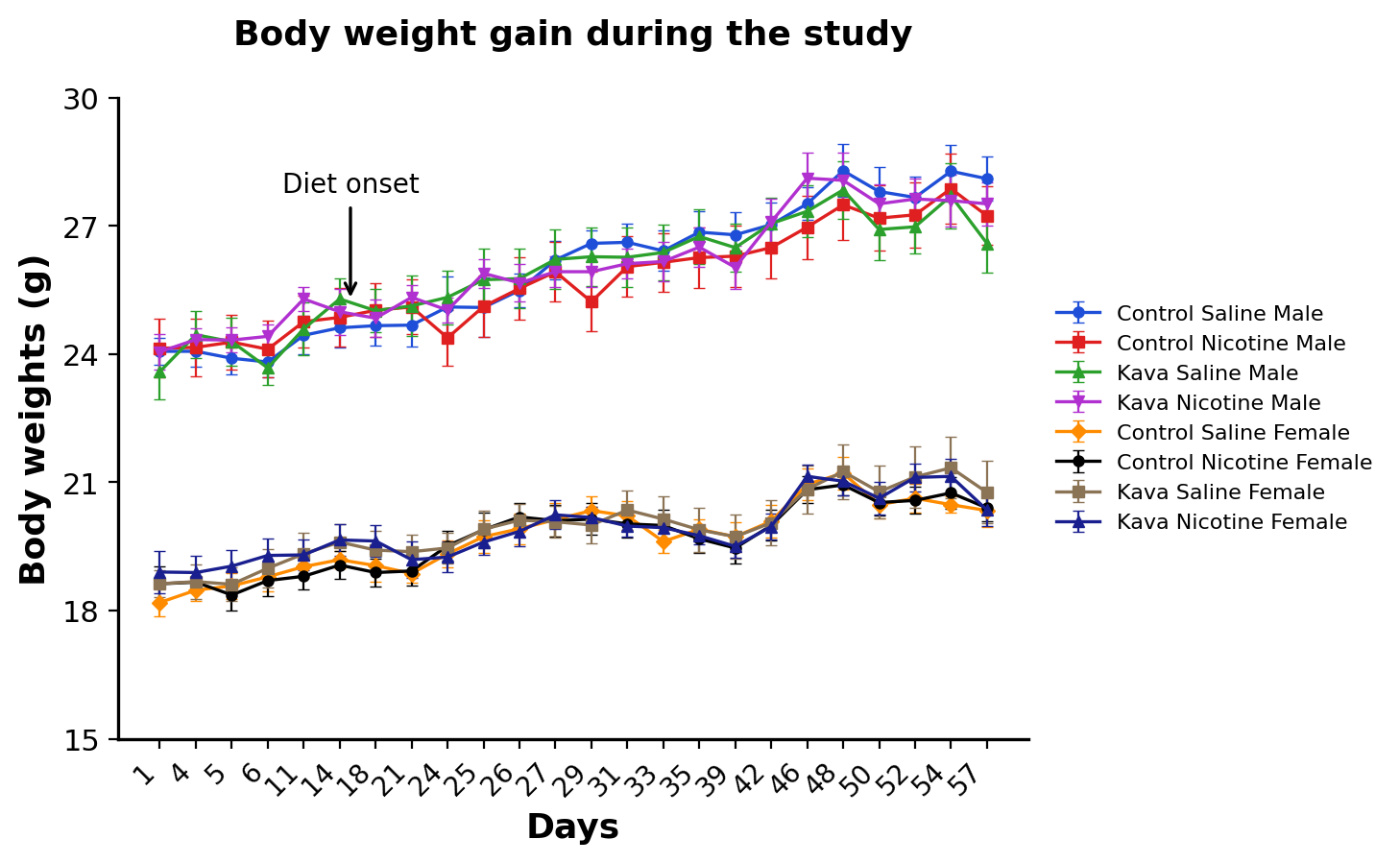
**Figure S2. Absolute body weight over time by sex, diet, and treatment during the study.** Body weight (g) was measured across 24 timepoints from Day 1 to Day 57 in male and female mice fed either a standard or kava-containing diet and treated with either saline or nicotine. The arrow indicates the start of the dietary intervention (Day 15); body weights prior to this point represent the pre-diet baseline period. Data are presented as mean ± SEM. N = 8–9 per group.

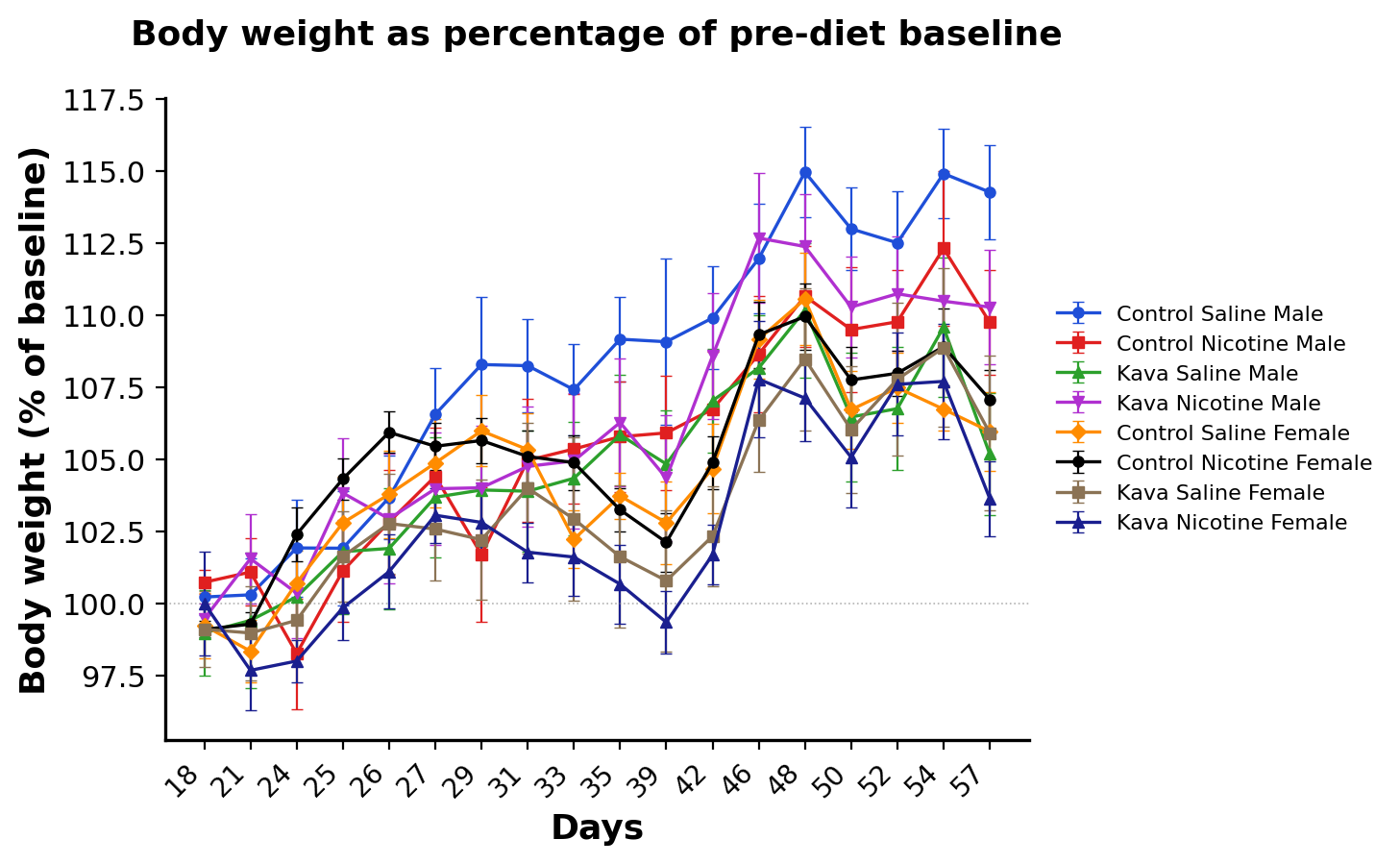
**Figure S3. Body weight expressed as a percentage of pre-diet baseline.** Body weight at each post-diet timepoint (Day 18–Day 57) was normalized to the body weight of each animal (Day 14 = 100%) to account for individual differences in starting weight. Data are shown for male and female mice fed either a control or kava-containing diet and treated with either saline or nicotine. The statistical analysis did not reveal a significant effect of diet or nicotine treatment on normalized body weight, while a significant effect of sex was observed, with males showing greater relative weight gain than females. Data are presented as mean ± SEM. N = 8–9 per group.
